# Modeling in an oral health study through two statistical methods in Uruguay

**DOI:** 10.1101/611921

**Authors:** Fernando Massa, Natalia Berberian, Ramón Álvarez-Vaz

**Affiliations:** Assistant Professor Universidad de la República, Facultad de Ciencias Económicas y de Administración, Departamento de Métodos Cuantitativos, Instituto de Estadística, Montevideo, Uruguay; Assistant Professor Universidad de la República, Facultad de Agronomía, Dpto. de Biometrìa, Estadística y Cómputo, Montevideo, Uruguay; Associated Professor Universidad de la República, Facultad de Ciencias Económicas y de Administración, Departamento de Métodos Cuantitativos, Instituto de Estadística, Montevideo, Uruguay

## Abstract

In epidemiological studies it is common practice to work with binary variables that reflect the presence of certain diseases, which in turn may be associated with another set of variables, that in general are assumed as risk factors of the former. In the field of epidemiological studies referred to oral health, it is common to inquire about the relationship between the presence of some pathologies and certain characteristics of the study participants through generalized linear models (GLM). However, this type of analysis is usually carried out for each variable of interest separately and at no time is a measure obtained that summarizes the status of each participant. The objective was to apply and compare two methodologies; one applying classical approach of explaining each oral disease separately from a set of explanatory variables and another using item response theory (IRT) models (specifically the Rasch model) since they allow the joint analysis of a set of variables obtaining an individual assessment as a by-product, which in this case is interpreted as “sickness proneness”. On the other hand, the analysis presented here extends the Rasch model including a linear predictor that allows to investigate about the possible effect of several factors on the propensity of the individuals to suffer the different pathologies. Our results found evidence of an effect of gender, insufficient physical activity (IPhA) and age on general proneness to oral diseases.

## Introduction

The epidemiological study of the most common oral pathologies, decay (D), loss of attachment (LoA), periodontal pockets (PP) and non functional dentition (NFD), can be carried out through different indicators, the most accepted of them is through binary variables representing the presence/absence of each pathology.

There is a hierarchy of the variables that will be used, those that appear in block one in Table 1 are risk factors, which are characterized as life habits. The variables in block two represent the most important noncommunicable diseases (NCD) and represent the largest burden of disease worldwide. Finally, the variables of block three represent oral disease, which can also be considered as NCD. Specialists think that these diseases are closely linked to the variables of both blocks one and two.

**Table 1.** Hierarchical blocks for Variables

| Block | Variable | Description |
| --- | --- | --- |
| 3*Behavioral | DS | Diary smoking |
|  | HAC | High alcohol consumption |
|  | IPhA | Insufficient physical activity |
| 4*Metabolic | BMI | Overweight / obesity |
|  | WHr | Waist / hip ratio |
|  | HBP | High blood pressure |
|  | Diab | Diabetes |
| 4*Oral Disease | PP | Periodontal pockets |
|  | NFD | Non functional dentition |
|  | D | Decay |
|  | LoA | Loss of attachment |

A very important characteristic to keep in mind that the variables of the three blocks are evaluated through quantitative variables, in order to determine if different risk factors were present or not, different thresholds were used, which allow to determine the presence off each risk factor. In the context of epidemiology, where NCD are studied, it is common practice to work with binary variables that reflect the presence of certain diseases, which in turn are associated with another set of diseases, called co-morbidities, also measured through variables binary and that in general are assumed as risk factors of the former. In the field of epidemiological studies, there are situations where NCD are managed, particularly in oral health, where both types of variables can be interchangeable as to who plays the role of risk factor.

In the field of health population surveys, it is a common practice for the epidemiological analysis investigate the factors that propitiate the occurrence of such pathologies using Generalized Linear Models. In this way it is possible to determine what are the conditions for a certain disease, however, these simple models are not able to carry out this analysis simultaneously in several pathologies. For this reason, we propose to use Item Response Theory (IRT) models [Baker,2017] in the epidemiological field because:

- they are capable of jointly analyze a set of outcomes and
- provide an assessment of each individual.

Some applications of this methodology to the dental area are [Reis,2014] on which the authors assess the psychometric properties of dental care questions to calculate a score for Brazilian states, [Yu-Feng,2013] used item response models to analyze an anesthesiology examination for medical and dental students and [Francisco,2012] who evaluate the oral health knowledge of a group of sixth grade students in a elementary school in the municipality of Araçatuba, São Paulo, Brazil.

However, the most frequently used IRT models provide indicators that describe the behavior of each variable without considering the possible effect of other set of explanatory variables. To overcome this difficulty we propose to model the outcomes through a Rasch model where the behavior of subject parameters is determined by a normal distribution whose mean is modeled by a linear predictor (see Fig) 1).

**Fig 1.**
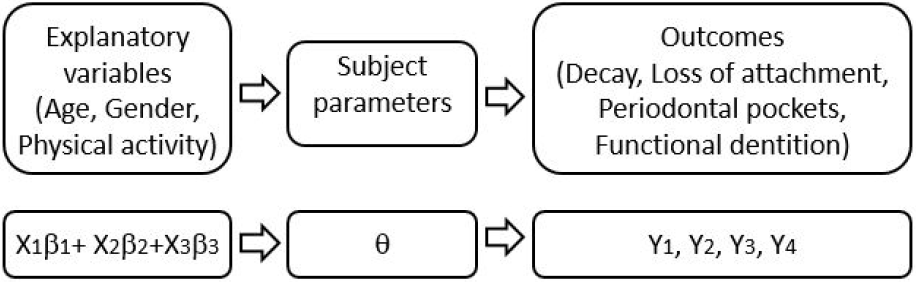
Flow diagram of the proposed model.

The data used comes from a study of people demanding attention in Dentistry Faculty, of the Universidad de la República, Montevideo, Uruguay, during the period 2015-2016. There were 602 participants where the presence/absence of four oral diseases was studied taking into account gender, age and physical activity of each participant.

## Materials and methods

As mentioned in the previous section, given the binary nature of the variables used for each oral disease, the Rasch model [Rasch,1960] appears as a natural starting point for the analysis.

The simplest IRT model assigns one “difficulty” parameter to every variable ([Baker, 2017]) and a non-observable parameter to each individual. Different theoretical curves from the Rasch model are shown in Figure 2, where *Y*_*ij*_ is the answer of individual *i* for the ‘item *j*.

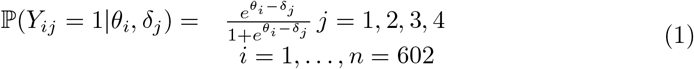

which is known as the Rasch model or one parameter logistic model.(see Fig 2)

**Fig 2.**
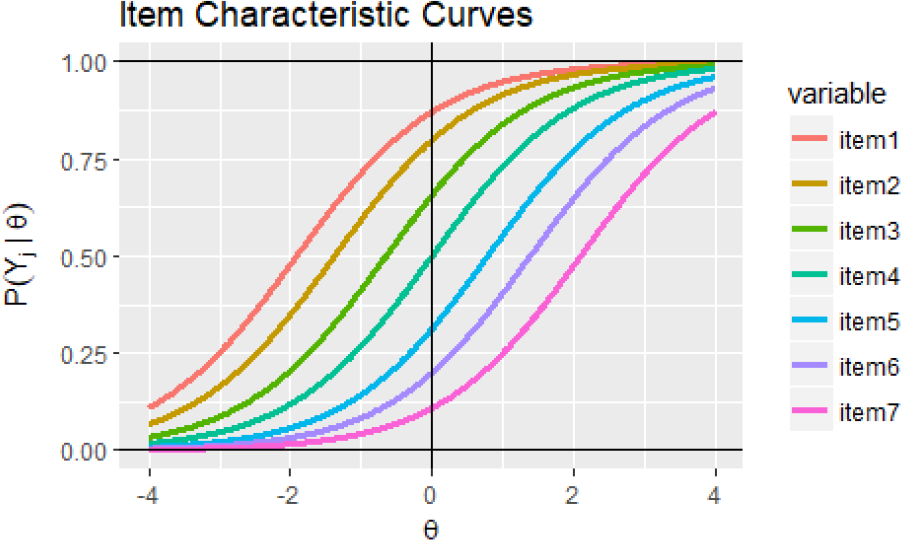
Different theoretical curves generated from a Rasch model.

We extended the model considering a set of predictors in the mean of the random effect used to model the behavior of each individual as is shown in equation 2. To compare our results with those obtained using classical analysis, we fitted a logistic regression model to the occurrence of each oral disease separately using the same set of predictor variables. Regarding the effect of explanatory variables, we considered a non-linear effect of age. The resulting curve was obtained using restricted cubic splines [Durrelman,1989] as part of the linear predictor. Finally, in order to get positive estimates for every risk factor, we included the variable Insufficient Physical Activity^1^ (IPhA)into the linear predictor instead of Physical activity.

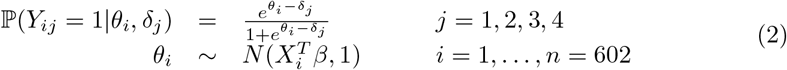

Where *Y*_*ij*_ represents the occurrence of the disease *j* on participant *i*, *θ*_*i*_ is the subject parameter (which in this context can be interpreted as the sickness proneness of each participant), *δ*_*j*_ is the difficulty parameter of each variable (that here is related to the prevalence of each pathology), 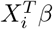 is a linear predictor that accounts for the effect of gender, age^2^ and physical activity. Equation 3 expresses the likelihood function.

The likelihood function of the model appears in the Equation 3

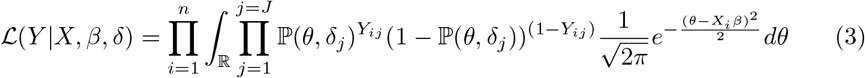

The optimization of the likelihood function is carried out numerically and, through it’s Hessian matrix, an approximation of the variance of each estimator is obtained. All calculations are carried out through the software R [RcoreTeam,2017].

## Results

A sample of 602 people who consult in the period corresponding to May 2015-June 2016 is applied, which are selected by systematic sampling, to which a sociodemographic questionnaire and a complete oral examination are applied, where the state of the teeth and mucosa is evaluated. From this information, the following attributes that make up 3 blocks of variables are considered for analysis and modeling.

**Table 2.** Prevalence of variables studied

| Block | Variable | Prevalence |
| --- | --- | --- |
| Behavioral | Diary Smoking | 0.331 |
|  | High alcohol consumption | 0.098 |
|  | Insufficient physical activity | 0.447 |
| Metabolic | Overweight / obesity | 0.573 |
|  | High Blood Pressure | 0.563 |
|  | Waist / hip ratio | 0.432 |
|  | Diabetes | 0.213 |
| Oral Disease | Periodontal pockets | 0.586 |
|  | Decay | 0.728 |
|  | Non Functional dentition | 0.596 |
|  | Loss of attachment | 0.636 |

The participants had an range age between 18 and 85 years old, having a mean of 45.021 (SD = 16.849) and there were 352 women and 250 men. As for the variables of the behavioral block, 199 smoked on a daily basis, 59 people had HAC and IPhA was detected on 269 participants. On the metabolic block, 345 participants presented overweight, 339 had HBP, 260 had an altered value of WHr and 128 had diabetes. As for the oral pathologies, the prevalence of PP,D and NFD and LoA were 0.586, 0.728, 0.403 and 0.636 respectively.

In addition to the risk factors considered, attributes such as age and gender were included in the model. The modeling strategy (after having estimated several models) is as follows:

1. Working with gender, a non-linear for age, the behavioral and metabolic risk factors in an IRT modeling framework.
2. Logistic Regression was estimated independently for each pathology using the same set of explanatory variables.

Joint analysis of four oral diseases and significant risk factors using the extended Rasch model is presented in Table 3.

**Table 3.**
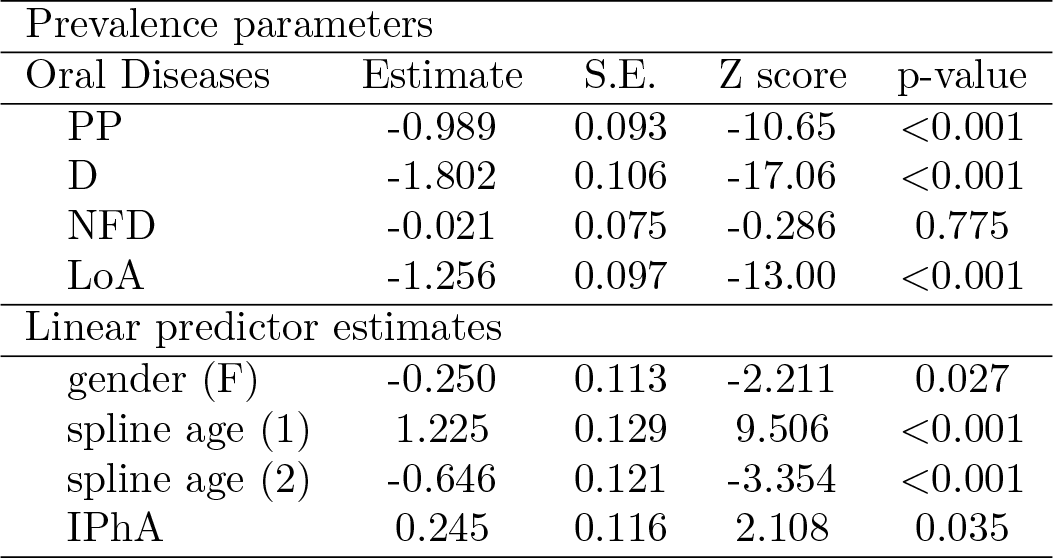
Estimates of the Rasch model with covariates

| Prevalence parameters |  |  |  |  |
| --- | --- | --- | --- | --- |
| Oral Diseases | Estimate | S.E. | Z score | p-value |
| PP | -0.989 | 0.093 | -10.65 | <0.001 |
| D | -1.802 | 0.106 | -17.06 | <0.001 |
| NFD | -0.021 | 0.075 | -0.286 | 0.775 |
| LoA | -1.256 | 0.097 | -13.00 | <0.001 |
| Linear predictor estimates |  |  |  |  |
| gender (F) | -0.250 | 0.113 | -2.211 | 0.027 |
| spline age (1) | 1.225 | 0.129 | 9.506 | <0.001 |
| spline age (2) | -0.646 | 0.121 | -3.354 | <0.001 |
| IPhA | 0.245 | 0.116 | 2.108 | 0.035 |

## Discussion

It can be observed that, because of the negative values of the difficulty parameters of the fitted Rasch model, the four pathologies have relatively high prevalence in the studied population. Figure 3 presents the expected value of “sickness proneness” as a non-linear function of age with different intercept for gender and physical activity status. It can be seen that men present higher proneness to oral diseases than women and, regarding insufficient physical activity, it decreases the mean value of “sickness proneness”, (see Fig 3).

Regarding the effect of age, the restricted cubic spline shows a significant increase effect (with negative concavity). Finally, a significant effect of physical activity, as well as gender, was found in the propensity to oral diseases.

**Fig 3.**
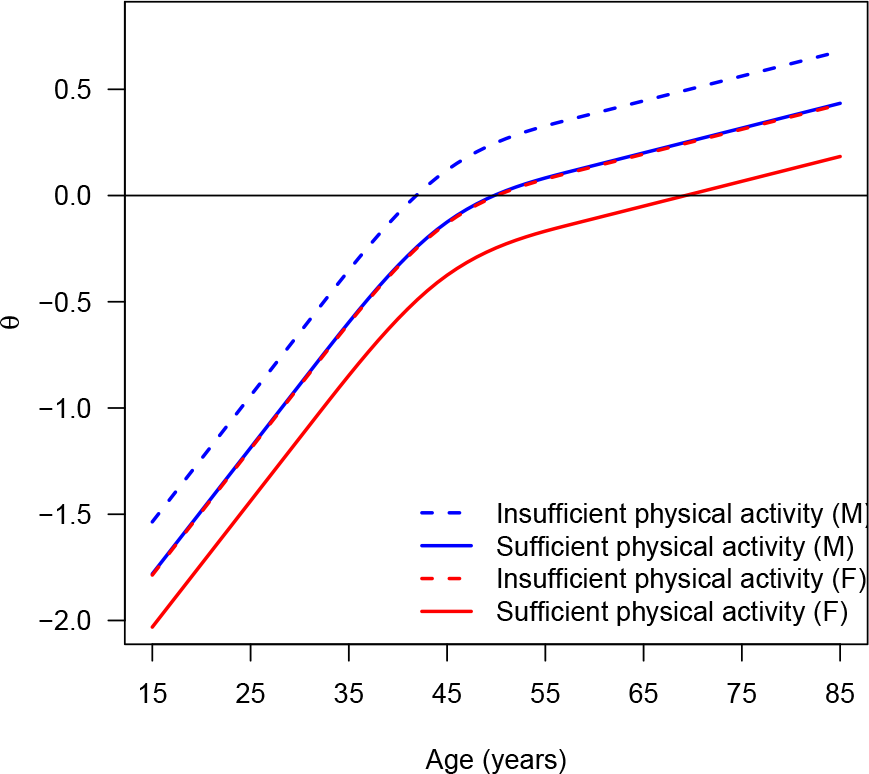
Sickness proneness according to age, gender and physical activity.

The results presented on Table 4 show the effect of age, gender and physical activity on every oral pathology separately. It can be seen that physical activity has a significant effect on the presence of decay as well as age is associated to every pathology.

**Table 4.** Logistic Regression Models.

|  | PP |  | D |  | LoA |  | NFD |  |
| --- | --- | --- | --- | --- | --- | --- | --- | --- |
| Variable | Estimate |  | Estimate |  | Estimate |  | Estimate |  |
| DS | 0,067 |  | 0,052 |  | 0,036 |  | 0,024 |  |
| HAC | 0,104 |  | -0,035 |  | 0,081 |  | -0,079 |  |
| IPhA | -0,003 |  | 0,097 | ** | 0,044 |  | 0,066 | . |
| BMI | 0,072 |  | 0,012 |  | 0,027 |  | 0,003 |  |
| WHr | -0,11 | . | 0,047 |  | -0,016 |  | 0,049 |  |
| HBP | 0,067 |  | 0,056 |  | -0,019 |  | -0,06 | ** |
| Diab | 0,042 |  | -0,081 |  | 0,06 |  | -0,018 |  |
| Age | 0,005 | ** | -0,004 | ** | 0,011 | ** | 0,016 | ** |
| Gender | -0,028 |  | -0,076 | . | -0,059 |  | 0,013 |  |

\*\* p-value ≤ 0.01 \* p-value ≤ 0.05 . p-value ≤ 0.10

It can be seen that, to a greater or a lesser extent, age can be related to every pathology,having a greater increase on early ages, slowing down as on late ages.In terms of gender, it can be seen that it has little or null effect on every pathology. Ultimately, IPhA shows a significant effect only on decay an non-functional dentition.

## Conclusion

Through the proposed model, it was possible to determine the prevalence level of the studied pathologies as well as the effect of some covariates of interest. It was observed that insufficient physical activity had a significant effect on “sickness proneness”.There was also detected a significant difference between men and women, while a non-linear (and increasing) effect of age was observed on the tendency to suffer from oral diseases. These results partially agreed with those of separated logistic regressions. The main difference between the two approaches was the effect of gender, while it was barely detected as significant on some oral pathologies, it was found that is was very significant on “sickness proneness”.

Future lines of research will include extending these our approach considering more elaborated IRT models. Consider more flexible IRT Models (2/3 parameters) and interpret their results in this context. By the other hand evaluate different cut-off thresholds for some of the covariables and their impact on the pathologies.

Also taking into account that there is a hierarchy in the variables of the 3 blocks, a multilevel analysis is proposed.

## Footnotes

1 According to thw World Health Organization, a person has IPhA if he/she practices less than 150 minutes of “moderated” physical activity in a week, or les than 75 minutes of “vigorous” physical activity in a week

2 The effect of age is modeled non-parametrically using restricted cubic splines.

